# Mutating different α-tubulin acetylation sites has distinct effects on axon terminal morphogenesis in *Drosophila melanogaster*

**DOI:** 10.1101/2025.10.27.684845

**Authors:** Helen Than, Chloe J. Welch, Ethan Schauer, Sophia Truijillo, Jill Wildonger

**Author notes:** Address correspondence to: Jill Wildonger.

## Abstract

Microtubules are created from uniform α- and β-tubulin building blocks but typically carry out a variety of specialized functions within a cell. The post-translational modification of tubulin is one means by which microtubule function can be tuned to match different cellular activities. While multiple sites of acetylation have been identified in tubulin, particularly α-tubulin, the effect of acetylation at different sites on microtubule function remains poorly characterized. Here, we took a genetic approach in *Drosophila* to disrupt three conserved sites of acetylation (K326, K370, K401) in endogenous α-tubulin and characterized the effects on neuronal development. Acetylation-blocking mutagenesis of α-tubulin K326 (K326A) perturbed larval locomotion and reduced axon terminal growth at the neuromuscular junction. These deficits were accompanied by a reduction in stable microtubules, suggesting that the α-tubulin K326A mutation exerts its effect by disrupting microtubule stability. In contrast, mutagenesis of α-tubulin K370 and K401 had virtually no effect on microtubule stability, suggesting that the effects of these mutations on axon terminal morphogenesis and survival may be mediated through a different mechanism. Altogether, the varied effects of these mutations suggests that acetylation at these three different sites may regulate different aspects of microtubule function within developing neurons.

## INTRODUCTION

Microtubules are an essential cytoskeletal component of many different cell types. Although microtubules are generated from a common pool of similar α- and β-tubulin subunits, they carry out a range of specialized processes, including propelling cell division, mediating intracellular transport, and providing structural support. In neurons and other cells, microtubule function is thought to be spatially and temporally regulated via the patterning of microtubules by a combination of different microtubule-associated proteins (MAPs) and the post-translational modification of tubulin (Ramkumar *et al*., 2018; Janke and Magiera, 2020; Iwanski and Kapitein, 2023; Teoh and Bartolini, 2025). Many post-translational modifications (PTMs), including polyglutamylation, polyglycylation, and detyrosination, occur on the C-terminal tails of α- and β-tubulin and have been well-studied. Another prominent PTM is acetylation. Nearly a dozen conserved acetylation sites have been identified in tubulin, particularly α-tubulin (Choudhary *et al*., 2009; Weinert *et al*., 2011; Lundby *et al*., 2012; Liu *et al*., 2015b; Hansen *et al*., 2019); however, only one of these sites, α-tubulin lysine 40 (K40), has been extensively studied.

Indeed, the acetylation of α-tubulin K40 has become synonymous with microtubule acetylation, despite the existence of these other acetylation sites. Recent studies from our lab have characterized a second conserved acetylation site in α-tubulin, K394, in fruit flies. We discovered that K394 acetylation regulates the development of the larval neuromuscular junction (NMJ) as well as an adult central brain structure called the mushroom body (Saunders *et al*., 2022; Welch *et al*., 2025). However, it remains largely unknown what role the acetylation of other α-tubulin residues may play in regulating microtubules, including whether these various modifications may selectively control distinct aspects of microtubule behavior and function.

To determine how tubulin acetylation may regulate microtubule function in cells *in vivo*, we took a mutagenesis approach in *Drosophila*. We took this approach as the enzymes that modify these sites are not known. Moreover, acetyltransferases and deacetylases often have multiple protein targets, making it difficult to pinpoint whether the effect of manipulating an enzyme is tubulin-related. Here, we focus on three α-tubulin sites: K326, K370, and K401. These sites have been reported to be acetylated across multiple organisms, from flies to mammals (Choudhary *et al*., 2009; Weinert *et al*., 2011; Lundby *et al*., 2012; Liu *et al*., 2015a; Liu *et al*., 2015b; Hansen *et al*., 2019), and our own mass spectrometry studies also indicate that these sites are acetylated in the axons of cultured rat neurons. Virtually nothing is known, however, regarding the biological significance of modifying these sites. These three α-tubulin residues reside in distinct locations: K326 is at the interface of the tubulin heterodimer, K370 faces the microtubule lumen (similar to the well-characterized K40), and K401 is positioned on the microtubule surface, where it can interact with MAPs. Thus, acetylation of these lysine residues has the potential to affect different aspects of microtubule function.

We investigated the effects of mutating endogenous α-tubulin K326, K370, and K401 on the development of the fly nervous system. Neurons have heavily modified microtubules, and the acetylation of α-tubulin K40 and K394 have been implicated in regulating microtubules in both developing and mature neurons (Janke and Magiera, 2020; Saunders *et al*., 2022; Teoh and Bartolini, 2025; Welch *et al*., 2025). The results of our current study reveal that mutagenesis of each of these three sites disrupts neuronal development in varying ways and to different degrees. These new data, combined with previous studies of α-tubulin K40 and K394, suggest that different patterns of α-tubulin acetylation may selectively tune different aspects of microtubule behavior (e.g., microtubule stability versus mediation of motor-based transport) within neurons and other cells.

## RESULTS AND DISCUSSION

### Effects of mutating α-tubulin acetylation sites K326, K370, and K401 on survival

Like other organisms, the fly genome has multiple α-tubulin genes that encode different α-tubulin proteins, called isotypes, that vary in their sequence and expression pattern. The *Drosophila melanogaster* genome contains four α-tubulin genes, whose names reflect their cytological location: *αTub84B*, *αTub84D*, *αTub64C*, and *αTub85E* (Sanchez *et al*., 1980; Kalfayan and Wensink, 1981; Mischke and Pardue, 1982). Our studies focus on *αTub84B*, which is the predominant α-tubulin gene in flies (Raff, 1984). αTub84B is broadly expressed, essential, and homologous to the major α-tubulin isotype in humans, TUBA1A (Figure 1A). In a previous study, we replaced the *αTub84B* gene with an *attP* site to facilitate the easy knock-in of new *αTub84B* alleles via site-mediated integration (Jenkins *et al*., 2017). Using this fly strain, we mutated three different conserved sites of acetylation (K326, K370, and K401) to alanine in endogenous *αTub84B* and subsequently assessed animal survival, behavior, and synaptic terminal development to determine the effects of blocking acetylation at these sites.

**FIGURE 1:**
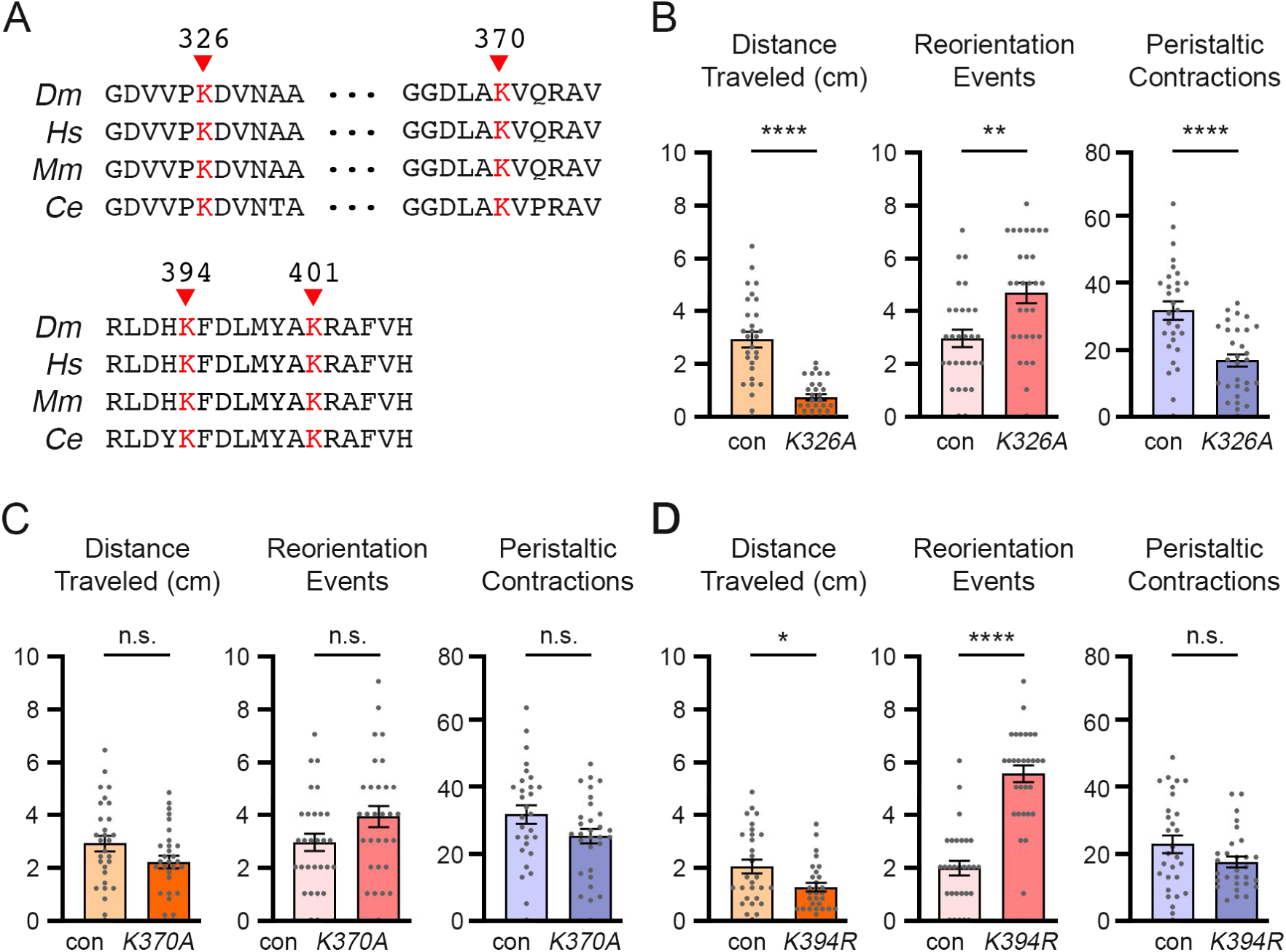
*αTub84B K326A* and *K394R* mutations disrupt larval locomotion. (A) Sequence alignment of acetylation sites in α-tubulin from flies (*Dm* αTub84B), humans (*Hs* TUBA1A), mice (*Mm* Tuba1a), and worms (*Ce* Tba-1). (B-D) Graphs showing the distance traveled (cm), number of peristaltic contractions, and number of reorientation events for control (*w^1118^*) and *K326A* (B), *K370* (C), and *K394R* (D) homozygous mutants. n = 29 (control) and 30 (*K326A*/*K326A*) (B); n = 29 (control) and 30 (*K370A*/*K370A*) (C); and n = 29 (control) and 29 (*K394R*/*K394R*) (D). The *K326A* and *K370A* experiments were performed in parallel, and the same control was used for each mutant. Statistical analysis: Two-tailed Student’s t-test with Welch’s correction (distance traveled, *K326A* and *K370A*; peristaltic contractions, *K370A* and *K394R*; reorientation events, *K326A*, *K370A*, and *K394R*) or Mann-Whitney U test (distance traveled, *K934R*; peristaltic contractions, *K326A*). n.s., not significant; *p<0.05; **p<0.01; ****p<0.000.

First, we asked whether these mutations affected animal survival. The *αTub84B K326A* and *K370A* mutants both survived to adulthood in approximately the expected proportions of homozygous and heterozygous animals, indicating that these two mutations do not affect viability (Table 1; 30% of the animals are expected to be homozygous). *αTub84B K326A* and *K370A* mutants are also viable in trans to an *αTub84B* deletion (*αTub84B^KO-attP^*). In contrast, the majority of *αTub84B K401A* homozygous mutant animals died during larval stages, with only a few surviving to pupal stages, indicating that the *K401A* mutation is lethal (Table 1). This lethality may be due to structural changes in α-tubulin caused by the substitution of alanine, a relatively small amino acid, for lysine. To test this possibility, we substituted K401 with arginine (K401R), an amino acid that is more similar to lysine in structure and charge (like alanine, however, arginine is not acetylated). We found that the *K401R* mutation also caused lethality and that homozygous mutant animals died at a similar developmental stage as the *K401A* mutants (Table 1; both *K401A* and *K401R* are lethal in trans to *αTub84B^KO-attP^*). Altogether, our results indicate that mutating αTub84B to block acetylation at K326 and K370 has no effect on viability whereas mutating K401 causes lethality, regardless of whether the substitution is conservative or not.

**TABLE 1.**
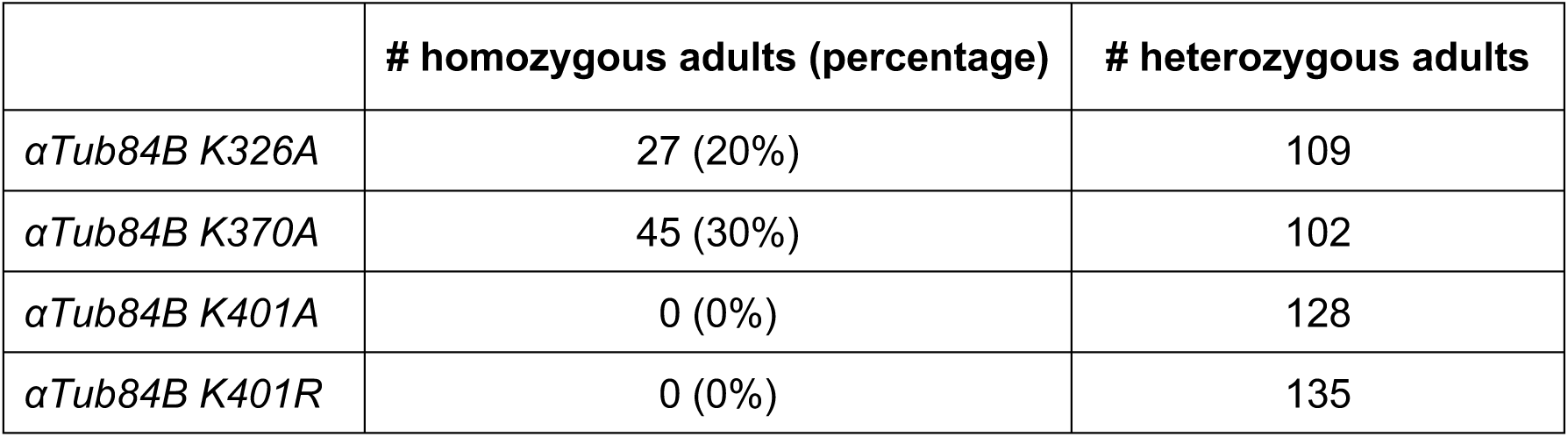
Effects of the *αTub84B* mutations on survival.

### The *αTub84B K326A* mutation, but not *K370A*, disrupts larval locomotion

Although the *αTub84B K326A* and *K370A* mutations do not affect survival, it is possible that they disrupt behavior. For example, mutating α-tubulin K40 does not affect survival, but the acetylation of K40 has a conserved role in mediating mechanosensation in flies and other organisms (Akella *et al*., 2010; Shida *et al*., 2010; Morley *et al*., 2016; Yan *et al*., 2018). Fruit fly larvae expressing αTub84B K40R are less responsive to mechanical stimuli and also display abnormal crawling behavior (Yan *et al*., 2018; Niu *et al*., 2023). To test the idea that the acetylation of α-tubulin K326 and K370 may be similarly involved in regulating behavior, we used a larval locomotion assay and quantified three parameters: distance traveled, number of reorientation events, and number of peristaltic contractions. We did not assay the *αTub84B K401A* and *K401R* mutants as insufficient numbers of larvae survive to the developmental stage at which the assay is performed. In addition to the *αTub84B K326A* and *K370A* mutants, we also evaluated an acetylation-blocking mutation of *K394* (*K394R*). We previously showed that the *αTub84B K394R* mutation disrupts the morphogenesis of motor neuron synaptic terminals, but we did not assay for any behavioral deficits, including disruptions in larval locomotion (Saunders *et al*., 2022).

We performed the larval locomotion assays and found that the *αTub84B K326A* mutation resulted in a significant reduction in both the distance-traveled and number of peristaltic contractions, which was coupled with an increase in the number of reorientation events (Figure 1B). In contrast, the *αTub84B K370A* mutation did not affect any of the larval locomotory parameters we assessed (Figure 1C). Similar to the *K326A* mutants, the *K394R* mutant animals traveled a significantly shorter distance than control animals and reorientated more frequently (Figure 1D). Since larval locomotion is regulated by motor neurons, the behavioral deficits displayed by the *K394R* mutants may reflect the changes in synaptic terminal morphogenesis at the NMJ that occur in the *K394R* mutant (Saunders *et al*., 2022). This raises the possibility that changes in synaptic terminal development may also occur in the *K326A* mutant animals.

### Mutations that disrupt the acetylation of αTub84B K326, K370, and K401 have differing effects on synaptic terminal morphogenesis and microtubule stability

To determine whether the acetylation of K326, as well as K370 and K401, might affect neuronal morphogenesis, we analyzed the growth of motoneuron axon terminals at the larval NMJ. The synaptic terminals of motoneuron axons are comprised of oval-shaped boutons, each of which houses multiple synapses, ranging from five to forty active zones (Menon *et al*., 2013; Aponte-Santiago and Littleton, 2020). Motoneuron axon terminals are divided into several types based on their physiological and morphological properties. We analyzed the glutamatergic type Ib ("big") and type Is ("small") axon terminals that innervate muscles 6 and 7 (m6/7). We visualized the axon terminals with anti-HRP, which labels neuronal membranes, and quantified the number of type Ib and type Is boutons. To identify stable microtubules at the NMJ, we used an antibody against Futsch, a microtubule-binding protein and MAP1B ortholog (Hummel *et al*., 2000; Roos *et al*., 2000). Futsch colocalizes with stable microtubules, which form loop-like structures in synaptic boutons; the number of these “Futsch loops” serves as a read-out of microtubule stability. We also probed for acetylation at α-tubulin K40 to determine whether this predominant acetyl mark is affected by disrupting other sites of acetylation in α-tubulin.

Our analysis of the *αTub84B K326A* mutant NMJs revealed a reduction in the overall number of synaptic boutons, reflecting a decrease in both type Ib and type Is boutons (Figure 2, A-C).

**FIGURE 2:**
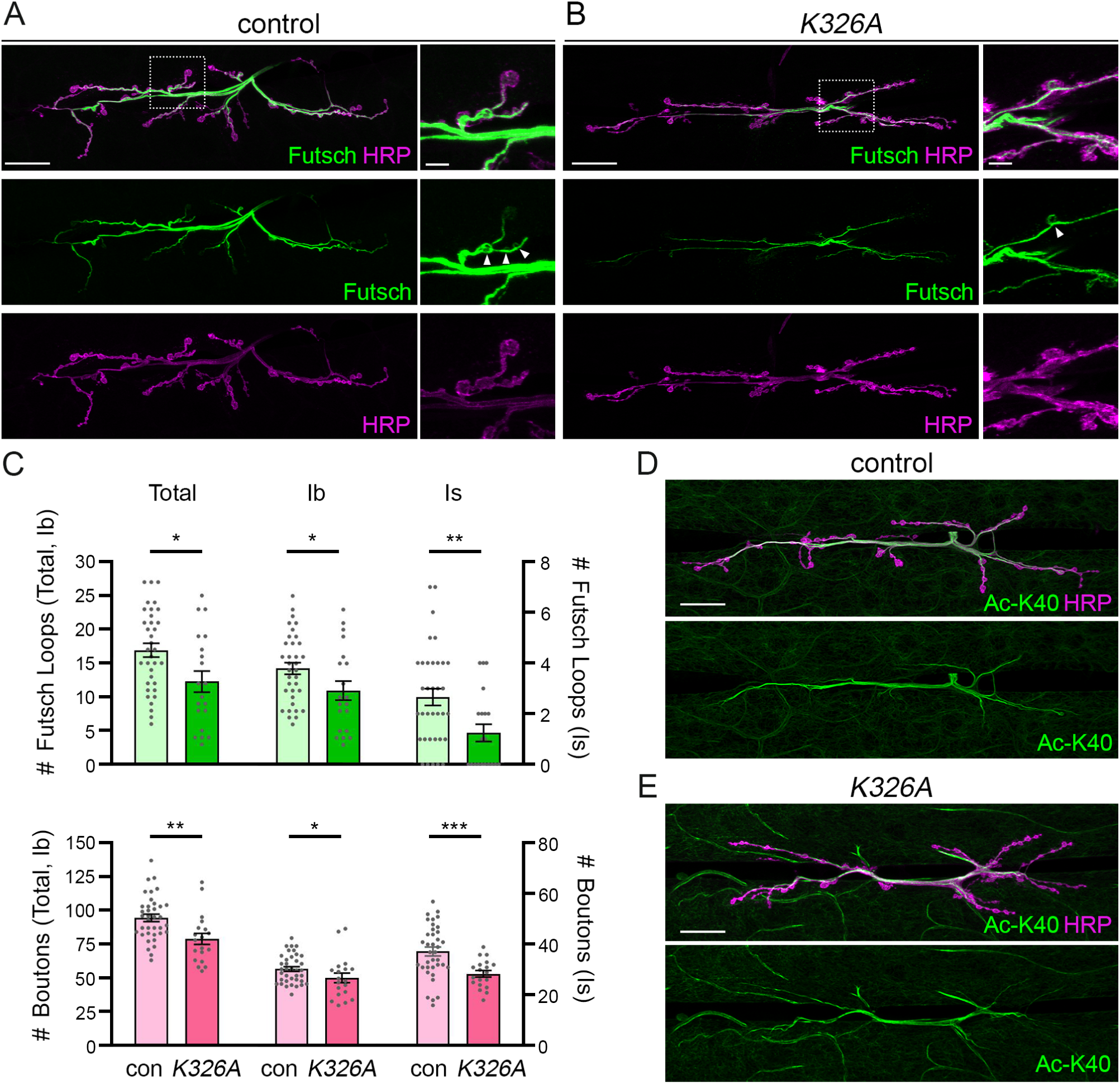
The numbers of boutons and Futsch loops are reduced at the developing NMJ in *αTub84B K326A* mutant larvae. (A and B) Images of type Ib and type Is axon terminals at m6/7 in control (*w^1118^*) (A) and *K326A*/*K326A* (B) third instar larvae. Neuronal membranes are labeled with HRP (magenta); stable neuronal microtubules are labeled with Futsch (green). Arrowheads indicate Futsch loops. (C) Quantification of Futsch loops (top) and boutons (bottom) in control and *K326A*/*K326A* mutant larvae. n = 37 (control) and 21 (*K326A*/*K326A*). Statistical analysis: Mann-Whitney U test. n.s., not significant; *p<0.05; **p<0.01; ***p<0.001. (D and E) Acetylation of α-tubulin K40 in control (*w^1118^*) (D) and *K326A*/*K326A* (E) larvae (magenta: HRP; green: anti-acetylated α-tubulin K40). Scale bars: 25 µm (left panels, A and B; D and E) and 5 µm (right panels, zoomed-in view, A and B).

Mirroring the reduction in bouton number, the number of Futsch loops was also reduced (Figure 2, A-C). There was no overt disruption in α-tubulin-K40 acetylation, indicating that this acetyl mark is not affected by the αTub84B K326A mutation (Figure 2, D and E). Thus, our results indicate that mutating αTub84B K326A disrupts both microtubule stability, as read-out by Futsch loops, and axon terminal morphogenesis. Given that perturbing NMJ development can affect larval motility, it is possible that the decrease in synaptic terminal boutons underlies the altered locomotory behavior of the *αTub84B K326A* mutant larvae.

While our analysis indicates that neuronal microtubule stability is reduced by the K326A mutation in endogenous α-tubulin, a study performed in HeLa cells found that exogenously expressed α-tubulin K326R has a modest but significant stabilizing effect on microtubules (Liu *et al*., 2015b). These differing results may reflect differences between endogenous versus exogenous mutant tubulin, or neurons *in vivo* versus immortalized cells in culture. α-tubulin K326 is positioned at the interface of tubulin heterodimers, where it is predicted to stabilize longitudinal "head-to-tail" interactions between the α-tubulin subunit of one dimer and the β-tubulin subunit of the neighboring dimer within a filament (Kumar *et al*., 2010; Chavez *et al*., 2019). A K326N mutation in human α-tubulin TUBA1A is associated with microlissencephaly, supporting the idea that K326 plays an important role in regulating microtubule structure and/or function (Kumar *et al*., 2010; Fallet-Bianco *et al*., 2014; Abbaali *et al*., 2023). Our findings are also consistent with the idea that mutating α-tubulin K326 alters microtubule stability.

While synaptic bouton number was reduced in the *αTub84B K326A* mutant, the *αTub84B K370A* mutant, in contrast, had significantly more boutons (Figure 3, A-C). Although the number of type Ib synaptic boutons increased, the number of Futsch loops in type Ib boutons was unchanged (Figure 3, A-C). In type Is boutons, the number of Futsch loops was reduced by approximately half. Similar to *αTub84B K326A*, the pattern and level of α-tubulin-K40 acetylation appeared normal (Figure 3, D and E). As the number of both type Ib and type Is boutons increased but Futsch was only affected in type Is boutons, a change in microtubule stability may not account for the increase in bouton number. Rather, it is possible that the K370A mutation may affect another aspect of microtubule function, such as transport, that might lead to the increase in bouton number. It is not clear why the *αTub84B K370A* mutation affects microtubule stability selectively in type Is boutons, although we previously observed type-specific effects on bouton growth in the *K394R* mutant (Saunders *et al*., 2022). The different physiological properties of, or other inherent differences between, the type Ib and type Is boutons may make them differentially sensitive to disruptions in the microtubule cytoskeleton (Kurdyak *et al*., 1994; Aponte-Santiago and Littleton, 2020). Although synaptic bouton numbers increased in the *αTub84B K370A* mutant, our results show that the *K370A* mutation has no effect on larval locomotion (Figure 1C). This indicates that the increase in bouton number is either not sufficient to cause a behavioral phenotype or is compensated for in some way.

**FIGURE 3:**
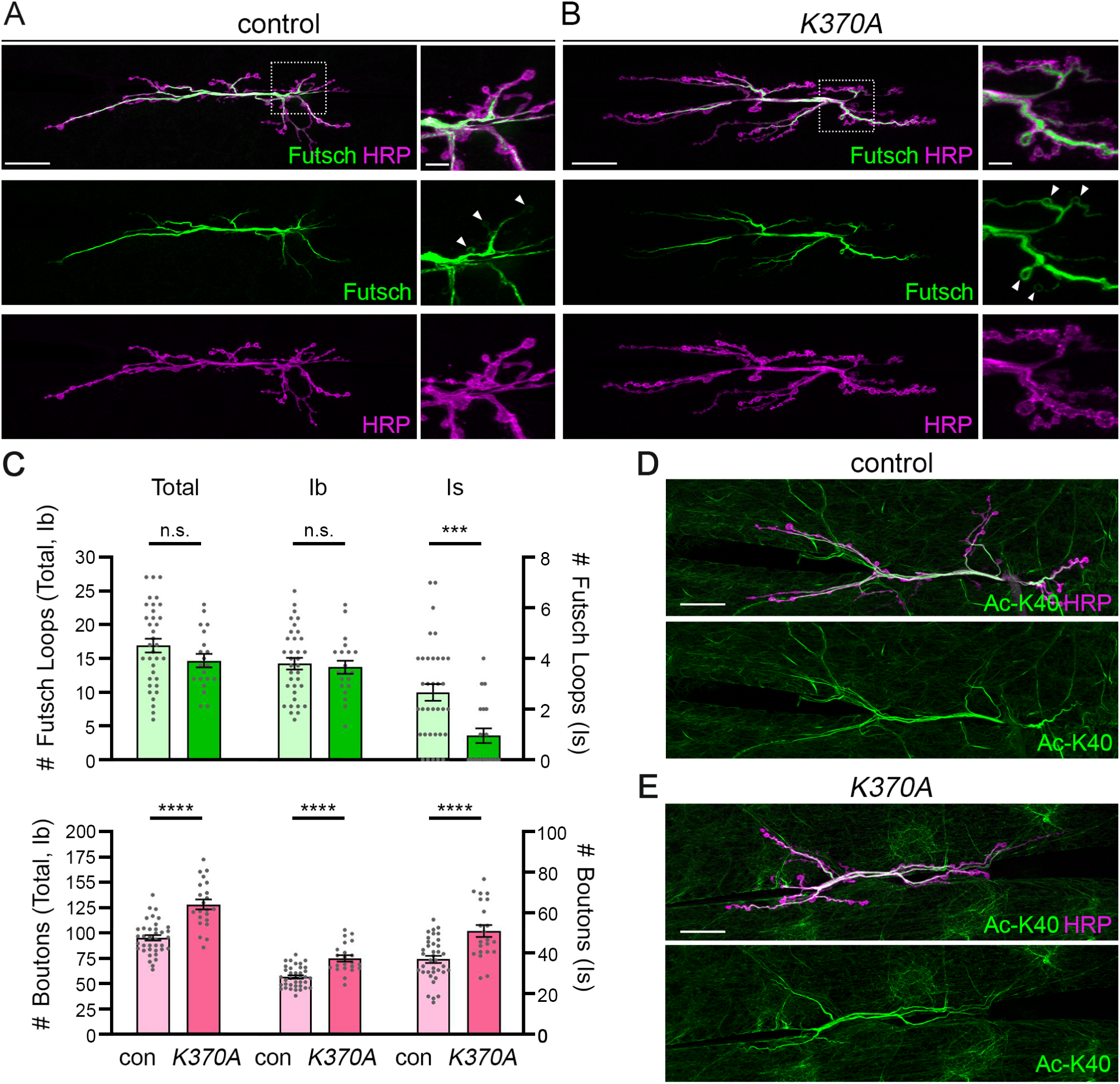
The *αTub84B K370A* mutation reduces bouton number and has a neuron-type specific effect on Futsch loops. (A and B) Images of type Ib and type Is axon terminals at m6/7 in control (*w^1118^*) (A) and *K370A*/*K370A* (B) third instar larvae. Neuronal membranes are labeled with HRP (magenta); stable neuronal microtubules are labeled with Futsch (green). Arrowheads indicate Futsch loops. (C) Quantification of Futsch loops (top) and boutons (bottom) in control and *K370A*/*K370A* mutant larvae. n = 37 (control) and 22 (*K370A*/*K370A*). Statistical analysis: Unpaired Student’s t-test and Mann Whitney U test (Is Futsch loops). n.s., not significant; ***p<0.001; ****p<0.0001. (D and E) α-tubulin K40 acetylation in control (*w^1118^*) (D) and *K370A*/*K370A* (E) larvae (magenta: HRP; green: acetylated α-tubulin K40). Scale bars: 25 µm (left panels, A and B; D and E) and 5 µm (right panels, zoomed-in view, A and B).

The *αTub84B K401A* and *K401R* mutations both resulted in lethality (Table 1). We decided to analyze the *αTub84B K401R* mutant, which has a conservative K-to-R mutation. The *αTub84B K401R* mutants displayed an increased number of type Ib boutons, but type Is bouton number was similar to controls, and the number of Futsch loops was unchanged (Figure 4, A-C). Like the *αTub84B K326* and *K370* mutants, the *αTub84B K401R* mutant displayed normal α-tubulin-K40 acetylation (Figure 4, D and E), indicating that K40 acetylation is not appreciably affected by disrupting acetylation at any of these three other sites in α-tubulin. The *K401R* mutation has a strong effect on animal survival, but, based on our findings, it is not clear what causes the animals to die. Similar to the *αTub84B K370* mutation, one possibility is that disrupting αTub84B K401 may affect the trafficking of a cargo(s) that is essential for neuronal communication and survival without creating any overt effect on microtubule stability. It is also possible that *αTub84B K401* perturbs a non-neuronal tissue or physiologic activity that is critical for survival.

**FIGURE 4:**
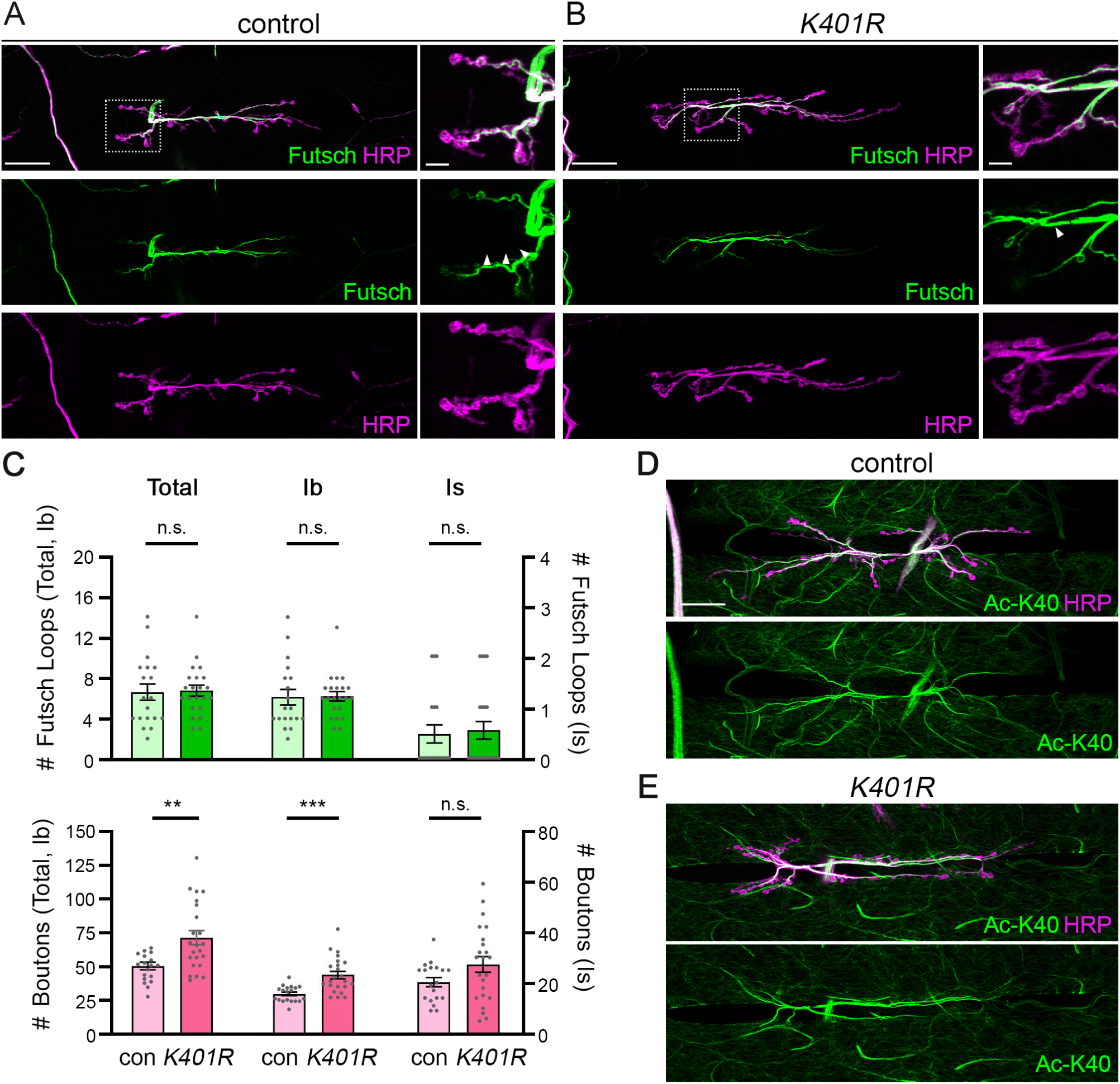
The *αTub84B K401R* mutation has a neuron-type specific effect on bouton number but does not affect Futsch loops. (A and B) Images of type Ib and type Is axon terminals at m6/7 in control (*w^1118^*) (A) and *K401R/K401R* (B) second instar larvae. Neuronal membranes are labeled with HRP (magenta); stable neuronal microtubules are labeled with Futsch (green). Arrowheads indicate Futsch loops. (C) Quantification of Futsch loops (top) and boutons (bottom) in control and *K401R/K401R* larvae. n = 19 (control) and 23 (*K401R/K401R*; boutons) and 22 (*K401R/K401R*; Futsch loops). Statistical analysis: Unpaired Student’s t-test (boutons, total Futsch loops) and Mann Whitney U test (type Ib and type Is Futsch loops). n.s., not significant; **p<0.01; ***p<0.001. (D and E) α-tubulin K40 acetylation in control (*w^1118^*) (D) and *K401R/K401R* (D) larvae (magenta: HRP; green: acetylated α-tubulin K40). Scale bars: 25 µm (left panels, A and B; D and E) and 5 µm (right panels, zoomed-in view, A and B).

In summary, our data reveal the effects of disrupting acetylation at α-tubulin K326, K370, and K401 via mutagenesis. Mutagenesis of these residues results in site-specific combinations of effects on survival, behavior, synaptic terminal morphogenesis, and microtubule stability. The variability in phenotypes suggests that these three sites and their modification may have different roles in regulating microtubules and microtubule-based activities. While we have focused on these residues as sites of acetylation, it is important to note that α-tubulin K326, K370, and K401 have been reported to undergo additional modifications. For example, all three sites have been reported to be ubiquitinated (Liu *et al*., 2015b). This raises the possibility that the effects of mutating a site may result from disrupting different, or multiple, modifications.

Isolating whether a phenotype is caused by perturbing one or several modifications will require identifying the relevant modifying enzymes. It is possible that the enzyme that acetylates α-tubulin K40, αTAT, also acetylates K370 given that both residues face the microtubule lumen. α-tubulin K326 and K401, however, do not face the microtubule lumen, making them unlikely targets of αTAT. Indeed, our previous study revealed that α-tubulin K394, which is positioned at the α- and β-tubulin interface within a dimer, is not acetylated by αTAT (Saunders *et al*., 2022). This suggests that there is another, currently unknown, acetyltransferase(s) that acts on α-tubulin. Similarly, the deacetylase HDAC6 targets α-tubulin K40, K394, and K370 but not K326 or K401 (Hubbert *et al*., 2002; Zhang *et al*., 2008; Liu *et al*., 2015a; Saunders *et al*., 2022).

Thus, it is likely that additional, and different combinations of, enzymes regulate the acetylation state of microtubules. Altogether, our studies point to the possibility that different α-tubulin sites are independently regulated, potentially to tailor different aspects of microtubule function with temporal or spatial control in cells.

## MATERIALS AND METHODS

### Fly husbandry and stocks

Fruit flies were maintained at 25°C on cornmeal-molasses-yeast medium. We obtained the control *w^1118^* strain from the Bloomington *Drosophila* Stock Center. The generation of new *αTub84B* alleles is described below. *αTub84B^KO-attP^* and *αTub84B K394R* are described in Jenkins et al., 2017 and Saunders et al. 2022, respectively.

### Creation of new *αTub84B* alleles

To create the new *αTub84B K326A*, *K370A*, *K401A*, and *K401R* alleles, the *pGE-attB-GMR* plasmid (DGRC Stock #1295) containing the new allele was injected into *αTub84B^KO-attP^* embryos expressing the integrase PhiC31 (BestGene Inc.) (Huang *et al*., 2009; Jenkins *et al*., 2017). The following mutations were introduced into *αTub84B* in *pGE-attB-GMR* using Phusion high-fidelity polymerase (New England Biolabs, M0530); the mutated sequence is underlined and in uppercase letters:

1. K326A: 5’-gatgttgtgccc<u>GCC</u>gacgtcaacgcc
2. K370A: 5’-ggtgatttggcc<u>GCC</u>gtgcagcgtgcc
3. K401A: 5’-ctgatgtacgcc<u>GCC</u>cgtgccttcgtc
4. K401R: 5’-ctgatgtacgcc<u>CGC</u>cgtgccttcgtc

### Fixation and Immunohistochemistry

Wandering second and third instar larvae were dissected in PHEM buffer (80 mM PIPES pH 6.9, 25 mM HEPES pH 7.0, 7 mM MgCl_2_, 1 mM EGTA) and fixed in 4% paraformaldehyde in 1X phosphate-buffered saline (PBS) with 3.2% sucrose for 30 min, permeabilized in 1XPBS with 0.3% Triton-X100 for 20 mins, quenched in 50 mM NH_4_Cl for 10 mins, blocked in buffer containing 2.5% BSA (catalog number A9647, Sigma), 0.25% fish skin gelatin (catalog number G7765, Sigma), 10 mM glycine, 50 mM NH_4_Cl, and 0.05% Triton-X100 for at least 1 h at room temperature. Fillets were then incubated in primary antibody in blocking buffer overnight at 4°C, washed in 1XPBS with 0.1% Triton-X100 at room temperature and incubated with secondary antibody in blocking buffer overnight at 4°C in the dark. After washing in 1XPBS with 0.1% Triton-X100, fillets were mounted in elvanol containing antifade (polyvinyl alcohol, Tris 8.5, glycerol, and DABCO; DABCO, catalog number 11247100, Fisher Scientific, Hampton, NH).

The following antibodies were used on dissected larval fillets: mouse anti-Futsch 22C10 (1:100, catalog number 22C10, Developmental Studies Hybridoma Bank, Iowa City, IA), mouse anti-acetylated-K40 6-11B-1 (1:500, catalog number T6793, Sigma-Aldrich), goat anti-HRP conjugated Alexa Fluor 647 (1:4000, or 0.5 μg mL-1, Jackson ImmunoResearch, West Grove, PA), Dylight 550 anti-mouse (1:2000, or 0.5 μg mL-1, ThermoFisher, Waltham, MA), Dylight 488 anti-mouse (1:1000, or 0.5 μg mL-1, ThermoFisher, Waltham, MA).

### Quantification of Larval Locomotion

Locomotor activity was measured using an adaptation of established methods (Nichols *et al*., 2012). Individual, wandering late third instar larvae were selected and placed at the center of a 100-centimeter (cm) Petri dish containing 2% agarose, which was set on top of a 0.2 cm^2^ grid. All larvae were allowed an acclimation period of 1 minute, followed by 1 minute of recording.

Videos were blindly scored to assess total distance traveled (number of grids crossed converted to cm), number of peristaltic contractions, and number of reorientation events.

### Quantification of Bouton Number and Futsch Loops

Type Ib and type Is boutons marked by anti-HRP and Futsch loops were quantified at m6/7 of segment A2 in blinded images. Boutons were counted in FIJI. Type Ib and type Is boutons were distinguished by size and anti-HRP intensity; along an axon terminal branch, type Is boutons are consistently smaller and have dimmer anti-HRP signal than type Ib boutons. Futsch loops were quantified as complete, unbroken loops of Futsch signal within a bouton.

### Quantification and statistical analysis

Data were blinded prior to analysis. Statistical analysis was performed in Excel and GraphPad Prism, using a significance level of p < 0.05. Data were first analyzed for normality using the D’Agostino-Pearson test or Shapiro-Wilk test. Normally distributed data were then analyzed for equal variance and significance using either an F-test and Student’s unpaired t-test (two samples). Data sets that were not normally distributed were analyzed using Mann-Whitney U test (two samples). Each graph and figure legend reports the exact n, which is the number of larvae (Figure 1) or NMJs (Figures 2-4) that were analyzed for the different experimental genotypes (each point in the graph represents an individual larva or NMJ). The statistical tests used to evaluate the data for each experiment are included in the figure legend. The error bars represent the standard error of the mean (s.e.m.); in the graph, the top of the bar indicates the mean.

## ACKNOWLEDGEMENTS

We thank current members of the Wildonger lab for their input, advice and comments on the project, and we thank past members of the Wildonger lab (Harriet Saunders, Dena Johnson-Schlitz, Brian Jenkins) for their contributions to early stages of this study. We acknowledge the Bloomington *Drosophila* Stock Center for fly stocks (NIH P40OD018537). The 22C10 anti-Futsch antibody was obtained from the Developmental Studies Hybridoma Bank, created by the NICHD of the NIH and maintained at The University of Iowa, Department of Biology, Iowa City, IA 52242. This work was supported by the National Institute of Neurological Disorders and Stroke, NIH (R01NS116373, J.W.); Pathways in Biological Sciences training grant, NIH (T32 GM133351, C.J.W); Ruth Stern Endowed Fellowship (C.J.W.); and a Curci PhD Fellowship, Curci Foundation (C.J.W.).

